# The CD4 T cell-independent IgG response during persistent virus infection favors emergence of neutralization-escape variants

**DOI:** 10.1101/2024.12.22.629980

**Authors:** Katelyn N Ayers, Matthew D Lauver, Kalynn M Alexander, Ge Jin, Kitt Paraiso, Alyssa Ochetto, Sonal Garg, Daniel J Goetschius, Susan L Hafenstein, Joseph Che-Yen Wang, Aron E Lukacher

## Abstract

How changes in the quality of anti-viral antibody (Ab) responses due to pre-existing or acquired CD4 T cell insufficiency affect virus evolution during persistent infection are unknown. Using mouse polyomavirus (MuPyV), we found that CD4 T cell depletion before infection results in short-lived plasma cells secreting low-avidity antiviral IgG with limited BCR diversity and weak virus-neutralizing ability. CD4 T cell deficiency during persistent infection incurs a shift from a T-dependent (TD) to T-independent (TI) Ab response, resembling the pre-existing TI Ab response. CD4 T cell loss before infection or during persistent infection is conducive for emergence of Ab-escape variants. Cryo-EM reconstruction of complexes of MuPyV virions with polyclonal IgG directly from infected mice with pre-existing or acquired CD4 T cell deficiency enabled visualization of shortfalls in TI IgG binding. By debilitating the antiviral IgG response, CD4 T cell deficiency sets the stage for outgrowth of variant viruses resistant to neutralization.

**ONE SENTENCE SUMMARY:** Pre-existing and acquired CD4 T cell deficiency facilitates outgrowth of Ab-escape viral variants during persistent infection.

## INTRODUCTION

Sustained humoral immunity is integral to defense against persistent viral infections. Evolution of viruses in their hosts can give rise to variants that prevent recognition by neutralizing Abs (nAbs). An antiviral nAb repertoire of limited diversity that targets viral capsids having few sites to neutralize infectivity can give rise to nAb-escape variants; i.e., therapy with anti-viral monoclonal Abs (mAbs) ^1–3^. Narrowed Ab repertoires can result from pre-existing immunity to parental virus, waning memory responses, or CD4 T cell immunodeficiency ^4–7^.

CD4 T cell-dependent (TD) Ab responses within germinal centers (GCs) drive immunoglobulin (Ig) class-switched recombination (CSR) and somatic hypermutation (SHM) to generate high-affinity Ab responses. GCs give rise to memory B cells and long-lived plasma cells (LLPCs) ^8^. Before expansion of antigen-specific CD4 T cells or under conditions of CD4 T cell deficiency, an innate-like CD4 T cell-independent (TI) Ab response may be engaged. This TI response produces predominantly of short-lived, IgM secreting plasma cells having germline Ig sequences. TI IgM responses are largely directed against bacteria. Accumulating literature, however, indicates that TI Ab responses may also undergo CSR and SMH in high inflammatory microenvironments ^9^. Influenza infection in mice lacking CD4 T cells elicits a protective IgG response, but of lower titer and shorter longevity than under TD conditions ^10^. Critically ill SARS-CoV-2 patients have fewer Tfh and GCs, yet mount a robust TI IgG response characterized by high Ab concentrations, affinity, and neutralization activity that fails to curb infection ^11^. Mouse polyomavirus (MuPyV) generates an IgG response in T cell-deficient mice that controls early infection ^12–14^. Whether this antiviral TI IgG response is maintained through persistent infection and has sufficient viral epitope diversity.

CD4 T cell deficiency, either inherited (e.g., idiopathic CD4 lymphopenia) or acquired (e.g., AIDS, monoclonal Ab (mAb) therapies), is a major condition predisposing to progressive multifocal leukoencephalopathy (PML), an aggressive brain disease caused by JC polyomavirus (PyV) ^15–18^. JCPyV-PML isolates frequently have nonsynonymous mutations in the PyV’s major capsid protein, VP1, that confer resistance to neutralization ^19–21^. These mutations are found in the four external loops of VP1 where the anti-PyV IgG epitopes reside ^22^. nAb-escape VP1 variants are similarly found in kidney transplant recipients with BKPyV-associated nephropathy ^23^. We recently demonstrated that nAb-escape VP1 mutations arise in B cell-deficient mice given an anti-VP1 mAb and CD4 T cell depleted ^1^.

Here, we asked whether TI conditions narrow the endogenous Ab repertoire and enable outgrowth of VP1 nAb escape variants. We developed two TI MuPyV infection models: (1) CD4 T cell depletion before infection to mimic “pre-existing” or inherited CD4 T cell deficiencies (e.g., idiopathic CD4 lymphopenia) and (2) CD4 T cell depletion during persistent infection to simulate “acquired” loss of CD4 T cells after infection with PyV (e.g., HIV-AIDS and PML-associated immunomodulatory therapies). In both models, CD4 T cell loss led to anti-MuPyV TI IgGs having weak avidity and lower B cell receptor (BCR) diversity than the TD Ab response.

Ab-secreting cells (ASCs) generated under TI conditions were short-lived. Cryo-EM reconstruction confirmed that IgGs under pre-existing and acquired TI conditions progressively lost the ability to bind virions. nAb-evading VP1 MuPyV variants emerged when sera from TI mice were serial passaged with MuPyV. In summary, the limited VP1 epitope coverage afforded by TI IgGs allows outgrowth of nAb-escape variants, an antecedent to human PyV diseases.

## RESULTS

### The neutralizing TI IgG response to MuPyV is low avidity

To evaluate pre-existing TI antiviral IgG response (**Fig. 1A**), wild-type (WT) mice were given a control IgG (TD mice) or a CD4-depleting mAb (TI mice) which impaired formation of GCs upon MuPyV infection (**Fig. 1B-C**). Sera from TI mice had MuPyV-specific, isotype-switched IgG, although at a lower concentration than TD mice (**Fig. 1D**; **Fig. S1C**). In contrast, IgM levels were equivalent between the TI and TD mice, peaking at 7 days post infection (dpi) but undetectable by 21 dpi (**Fig. 1D**). Numbers of MuPyV-specific Ab-secreting cells (ASCs) were reduced in the spleen, bone marrow (BM), and kidney of TI mice compared to TD mice at 21 dpi with dramatically lower levels at 128 dpi (**Fig. 1E**). Antiviral TI IgGs, unlike TD IgGs, exhibited no increase in avidity towards MuPyV over the course of persistent infection (**Fig. 1F**). These findings indicate that anti-MuPyV TI Abs undergo CSR but fail to increase in magnitude and avidity during persistent infection.

**Figure 1.**
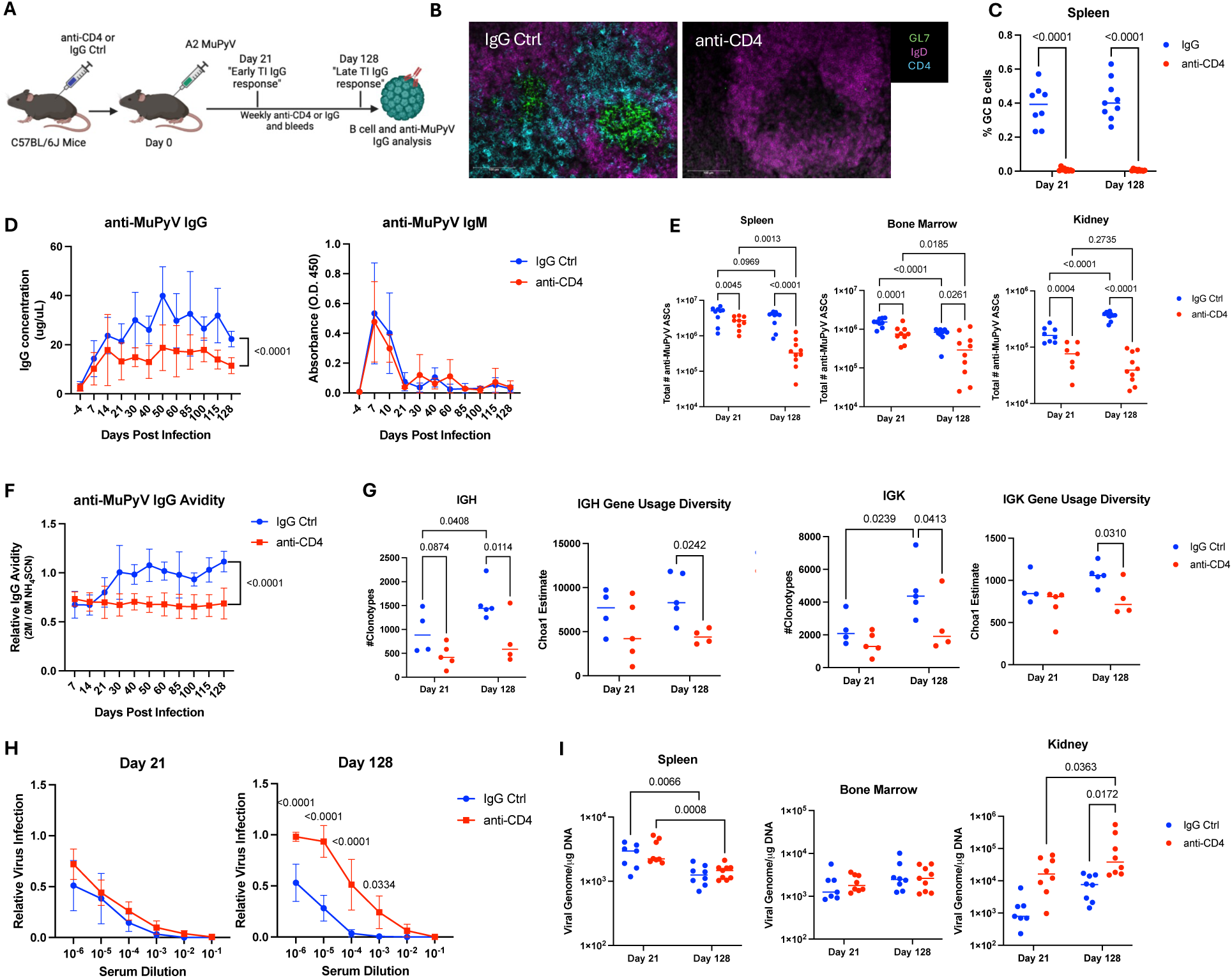
The TI Ab response to MuPyV infection is characterized by low avidity, virus-neutralizing IgG. **(A)** Experimental setup. B6 mice received depleting CD4 mAb (clone GK1.5) at 4- and 1-day pre-inoculation s.c. with MuPyV and then weekly post infection. Image made with Biorender. **(B, left)** Representative immunofluorescence staining of GCs - GL7 (green), IgD (magenta), and CD4 (cyan) in the spleen of TD and TI mice at 30 dpi. **(C)** Frequency of GC B cells (B220^+^ CD19^+^ IgD^-^ GL7^+^ CD95^+^) in the spleen at 21 and 128 dpi (n = 8). **(D)** Anti-MuPyV IgG and IgM in the sera of control and CD4 T cell-depleted mice as quantified by ELISA (n =10-15). **(E)** MuPyV-specific ASCs from spleen, BM, and kidneys were quantified by ELISpot assays (n = 8-9). The number of MuPyV-specific ASCs was calculated for the entire organ. **(F)** Anti-IgG avidity in CD4 T cell-depleted and control mice determined by ELISA with the addition of 2M NH_4_SCN (n = 10-15). **(G)** BCR sequencing of mRNA from FACS-sorted CD19^+^IgD^-^ splenic B cells from TI and TD mice at 21- and 128 dpi (n = 4-5). **(H)** Neutralization of purified MuPyV with serial dilutions of sera from CD4 T cell-depleted and control mice at 21- and 128 dpi (n = 8). **(I)** Viral DNA levels in BM, spleen, and kidney at 21 and 128 dpi (n = 8-9). Data are from 2-3 independent experiments. Data was analyzed by Two-way ANOVA with Tukey’s multiple comparisons test (**C, E, G, I**); XY analysis non-linear regression fit with extra- sum-of-squares F test (**D, F**); and two-way ANOVA with Šídák’s multiple comparison test (**H**).

BCR sequencing on activated B cells (IgD^-^ CD19^+^) revealed a narrowing of diversity under TI conditions. At 21 dpi, the number of clonotypes and BCR diversity was comparable between TI and TD mice B cells for both the heavy (IGH) and light (IGK) chains. At 128 dpi, however, TI BCR diversity was significantly lower than for TD B cells (**Fig. 1G**). Comparing early versus late Ab responses, BCR diversity increased in the TD B cells over time but not in TI B cells (**Fig. 1G**). Together, these data show that although CSR occurs in the TI mice, the TI IgG repertoire is constrained by a lack of SHM.

We next asked if the TI vs. TD IgGs differed in controlling MuPyV infection. At 21 dpi, sera from TI and TD mouse sera exhibited equivalent neutralization efficiency (**Fig.1H**). As infection progressed, TI Abs showed reduced ability to neutralize MuPyV than TD Abs (**Fig. 1H**). Despite these variations, viral titers in the spleen and BM at either time were equivalent between the TI mice and TD mice (**Fig. 1I**). Notably, kidney viral titers increased by 128 dpi (**Fig. 1I**), aligning with our previous work showing that CD4 T cell deficiency results in viral resurgence in the kidney ^1^.

To exclude the possibility that lymphopenia created by CD4 T cell depletion affect the MuPyV-specific TI IgG response, we examined the TI IgG response in MHCII KO mice, IL-21R KO mice, or CD40L blockade (**Fig. S2A**) ^24–26^. Each of these TI models recapitulated the findings with CD4 mAb-mediated depletion, including lack of GCs (**Fig. S2B**) and MuPyV- specific IgG response of lower avidity (**Fig. S2C-D**, **S2F-G**). Sera from 20 dpi, but not 100 dpi, neutralized MuPyV (**Sup.** Fig. 2E, H). Thus, mice with TI conditions mount a neutralizing IgG response comparable to healthy mice during early stages of viral infection, but during late persistent infection the TI IgGs are of lower titer, avidity, and BCR diversity than TD IgGs.

**Figure 2.**
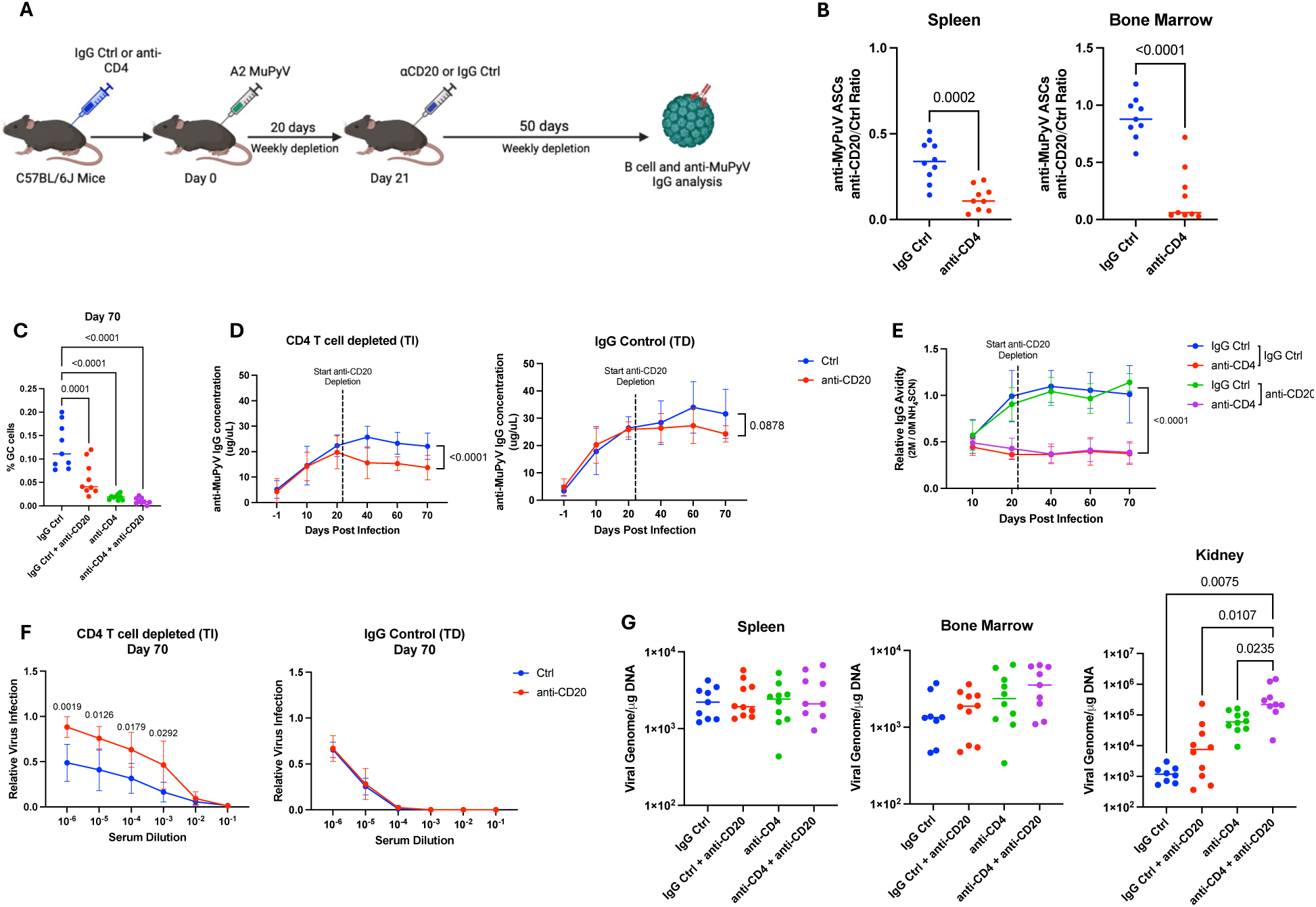
B cell depletion results in fewer anti-MuPyV IgG producing ASCs in CD4 T cell-deficient mice. **(A)** Experimental setup. CD4 depleted and IgG control B6 mice (Fig 1A) were given control IgG or CD20-depleting mAb starting at 20 and at 22 dpi mice were and then weekly until 70 dpi. Image made with Biorender**. (B)** Ratio of CD20^+^ B cell-depleted to control mice ASCs in the (left) BM and (right) spleen of CD4 T cell-depleted and CD4 T cell-competent mice (n = 9-10). **(C)** Frequency of GC B cells in the spleen at 70 dpi by flow cytometry. **(D)** Anti-MuPyV IgG concentration overtime in sera of anti-CD4 or IgG control mice with and without CD20-depleting mAb (n = 8). **(E)** Avidity of MuPyV-specific IgGs in the serum (n = 8- 9). **(F)** MuPyV neutralization by serum from anti-CD4 treated or IgG control mice with and without CD20-depleting mAb (n = 9-10). (**G**) Viral DNA levels in BM, spleen, and kidney at 70 dpi. (n = 8-9). Data are from 2-3 independent experiments. Data was analyzed by Student’s *t*-test **(B)**, XY analysis non-linear regression fit with extra-sum-of-squares F test **(D-E)**, or two-way Anova with Tukey’s multiple comparison test **(C, G);** two-way ANOVA with Šídák’s multiple comparison test (**F**).

### B cell depletion under TI conditions results in fewer anti-MuPyV IgG-producing ASCs

Because we detected TI MuPyV-specific ASCs at 128 dpi (**Fig. 1C-D**), we asked whether the TI ASCs were LLPCs. WT mice were CD4 T cell-depleted prior to infection and then given a CD20-depleting mAb starting 21 dpi to generate an anti-MuPyV ASC population. CD20 mAb depletion eliminates B cells but leaves ASCs intact ^27^. The ratio of ASCs in the spleen and BM of TI mice given anti-CD20 mAb vs. non-depleted control was significantly lower than the ratio of ASCs in the TD mice (**Fig. 2B**). Notably, with anti-CD20 the ratio of ASCs in the BM was higher than in the spleen of TD mice, suggesting that LLPCs are maintained in the BM but generation of new ASCs is disrupted in the spleen (**Fig. 2B**). Concurrently, GC B cell numbers were significantly reduced in the anti-CD20 treated TD mice, correlating with the loss of ASCs in the spleen (**Fig. 2C**).

In line with having fewer ASCs, TI mice had significantly lower IgG levels after CD20 B cell depletion. Conversely, no significant difference in the IgG concentration was seen in TD mice given CD20-depleting mAb (**Fig. 2D**). CD20-depleted TD mice maintained high avidity virus-specific IgG (**Fig. 2E**). In contrast, TI mice had low avidity anti-MuPyV IgG regardless of CD20 B cell depletion (**Fig. 2E**). TI IgGs also had significantly reduced virus neutralization activity after CD20 depletion (**Fig. 2F**). Virus levels were similar in the spleen and BM of CD20 B cell-deficient and -sufficient mice (**Fig. 2G**). Combined CD4 T cell- and CD20 B cell-depletion resulted in significantly higher virus levels in the kidney (**Fig. 2G**). Taken together, these data show the TI ASCs towards MuPyV are short-lived and need to be continuously replenished to maintain a strongly neutralizing TI IgG response to persistent MuPyV infection.

### Acquired CD4 T cell deficiency dampens the anti-MuPyV IgG response

MuPyV-infected WT mice received CD4 T cell-depleting mAb or rat IgG control beginning at 28 dpi (**Fig. 3A, S1B**). Acquired TI mice had fewer GCs, smaller GC area, and fewer GC B cells by 60 dpi (**Fig 3B-**D). No GCs were observed in spleens of acquired TI mice at 200 dpi (**Fig. 3B, 3D**), indicating that maintenance of GCs requires CD4 T cells. Acquired TI mice had fewer ASCs than TD mice at both 60 and 200 dpi in the spleen. The BM, however, had comparable numbers of MuPyV-ASCs at 60 dpi, but TI mice had lower numbers of ASCs than the TD mice at 200 dpi (**Fig 3E**). By extension, anti-MuPyV IgG titer and avidity fell significantly with acquired TI (**Figs. 3F-G**). Acquired TI and TD mice sera neutralized MuPyV equivalently at 60 dpi (**Fig. 3H**), but the TI sera poorly neutralized MuPyV by 200 dpi (**Fig. 3H**). Acquired TI and TD mice had equivalent viral titers in the spleen and BM at both 60 and 200 dpi; however, viral titers in the kidney increased from 60 to 200 dpi in the acquired TI mice like the pre-existing TI mice (**Fig. 3I**, **Fig. 1H**). These findings were confirmed in persistently infected mice given a CD40L blocking mAb at 28 dpi (**Fig. S3A**) which resulted in the loss of GCs (**Fig. S3B-C**), decreased number of ASCs in the spleen and BM (**Fig. S3D**), reduced anti-MuPyV IgG titers, and avidity (**Fig. S3E-F**), and neutralization efficacy at 200 dpi (**Fig. S3G**). Virus was controlled in the spleen and BM, as well as the kidney (**Fig. S3H**), suggesting that loss of CD4 T cells is required for virus resurgence in the kidney. These data indicate that acquired CD4 T cell deficiency incurs a shift from a TD to TI IgG response that parallels the lower titer, avidity, and neutralizing efficacy of the pre-existing TI anti-MuPyV IgG response.

**Figure 3.**
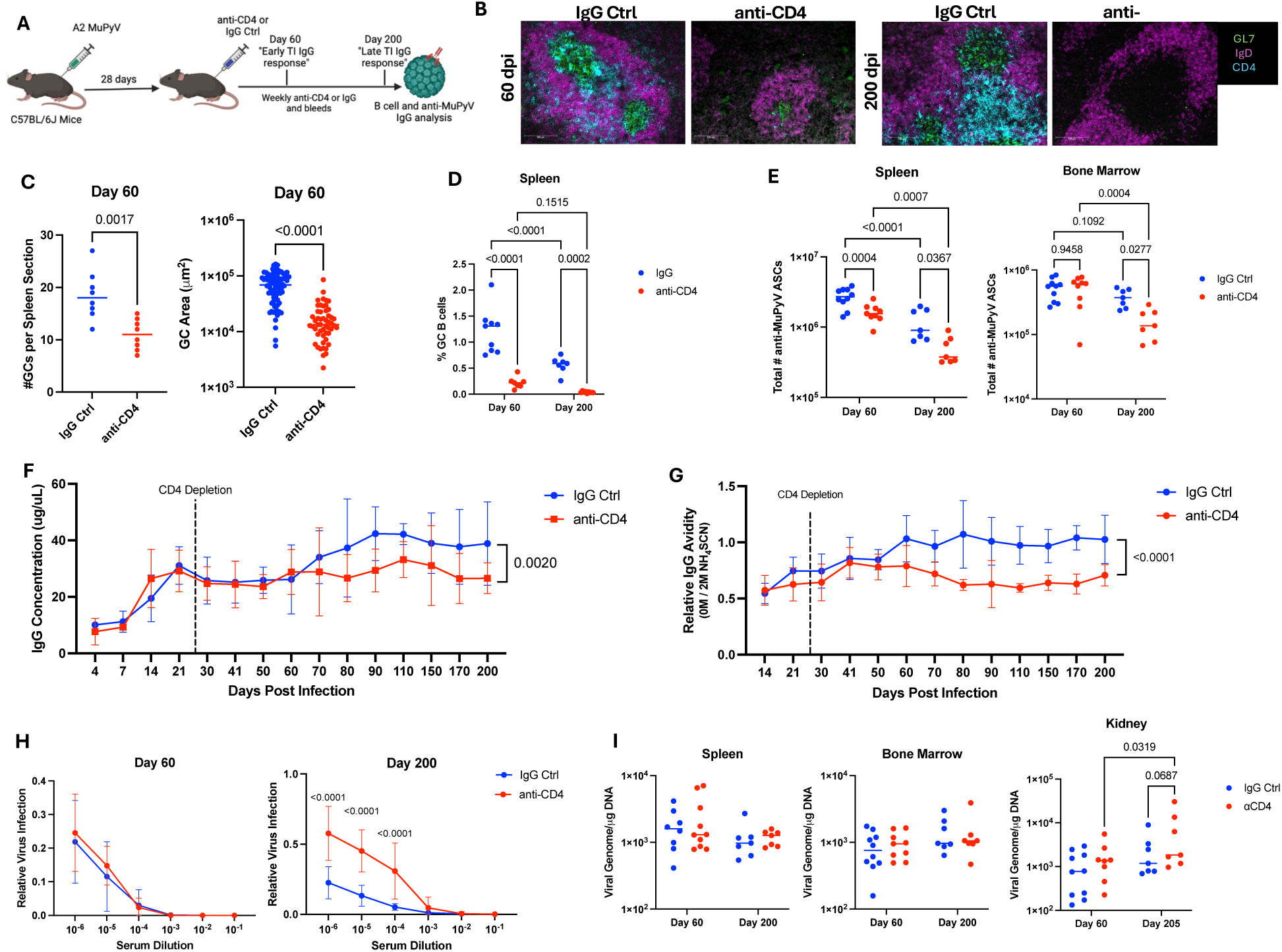
Acquired D4 T cell deficiency approximates the pre-existing CD4 T cell deficient TI anti-viral IgG response. **(A)** Experimental setup. B6 mice inoculated s.c. with MuPyV received CD4-depleting mAb or a control IgG weekly starting at 28- and 30- dpi. thereafter. Terminal samples were taken at 60 or 200 dpi. Image made with Biorender**. (B)** Representative immunofluorescence images of GCs [GL7 (green), IgD (magenta), and CD4 (cyan)] in the spleen of CD4 T cell-depleted or IgG control mice (n = 4). **(C, left)** Number of GCs in each spleen section at 60 dpi. GC were determined by GL7 bordered by IgD. **(C, right)** The area of the GCs in the spleen of acquired CD4 T cell deficient mice compared to IgG control mice at 60 dpi. **(D)** Frequency of IgD^-^ CD19^+^ GL7^+^ CD95^+^ GC B cells (n = 9-10). **(E)** Total number of MuPyV-specific ASCs in the **(left)** BM or **(right)** spleen cells. **(F)** Concentration and **(G)** avidity of MuPyV-specific IgG in the sera of CD4-depleted or IgG control mice by ELISA (n = 12). **(H)** Neutralization of MuPyV by sera from CD4 T cell-depleted or IgG control mice at **(left)** 60 and **(right)** 200 dpi (n = 8). **(I)** Viral DNA levels in the BM, spleen, and kidney at 60 and 200 dpi (n = 7-9). Data are from 2-3 independent experiments. Data was analyzed by Student’s *t*-test (**C**); two-way ANOVA with Tukey’s multiple comparisons test (**D, E, I**); XY analysis non-linear regression fit with extra-sum-of-squares F test (**F, G**); and two-way ANOVA with Šídák’s multiple comparison test (**H**).

### TI IgG fail to prevent emergence of nAb-evading, VP1 variant virus

MuPyV was passaged in the presence of serum from TI and TD mice. nAb-escape viruses were sequenced for VP1 mutations. No virus was detected following serial passaging with serum from either TI or TD mice up to 55 dpi; but virus plaques were first seen serial passage serum from TI mice at 85 dpi (**Table 1**). No mutations in VP1 were found in virus isolated from plaques using sera from mice at 85 dpi, 105 dpi, and one of the 285 dpi mice (**Table 1**). Serial passaging with serum from a TI mouse at 285 dpi, however, yielded a virus with a glutamic acid (E)-to-glycine (G) point mutation at residue 91 in the BC loop of VP1 of MuPyV (**Table 1**, **Fig. 4A**) ^28,29^. This E91G mutation (GAA) differs from the mutation introduced by site directed mutagenesis (GAG) in prior publications ^29^. Sera from both TI and TD mice recognized MuPyV.E91G (**Fig. 4B**) but exhibited lower avidity (**Fig. 4C**) and neutralization efficacy (**Fig. 4D**) against this VP1 mutation. We also detected the VP1 mutation E-to-lysine (K) at residue 68 in sera from one of three mice with acquired TI at 200 dpi (**Table 1**). These data fit the idea that TI IgGs lose avidity during persistent infection, allowing breakthrough of WT virus and the eventual emergence of variants with nAb-resistant VP1 mutations.

**Figure 4.**
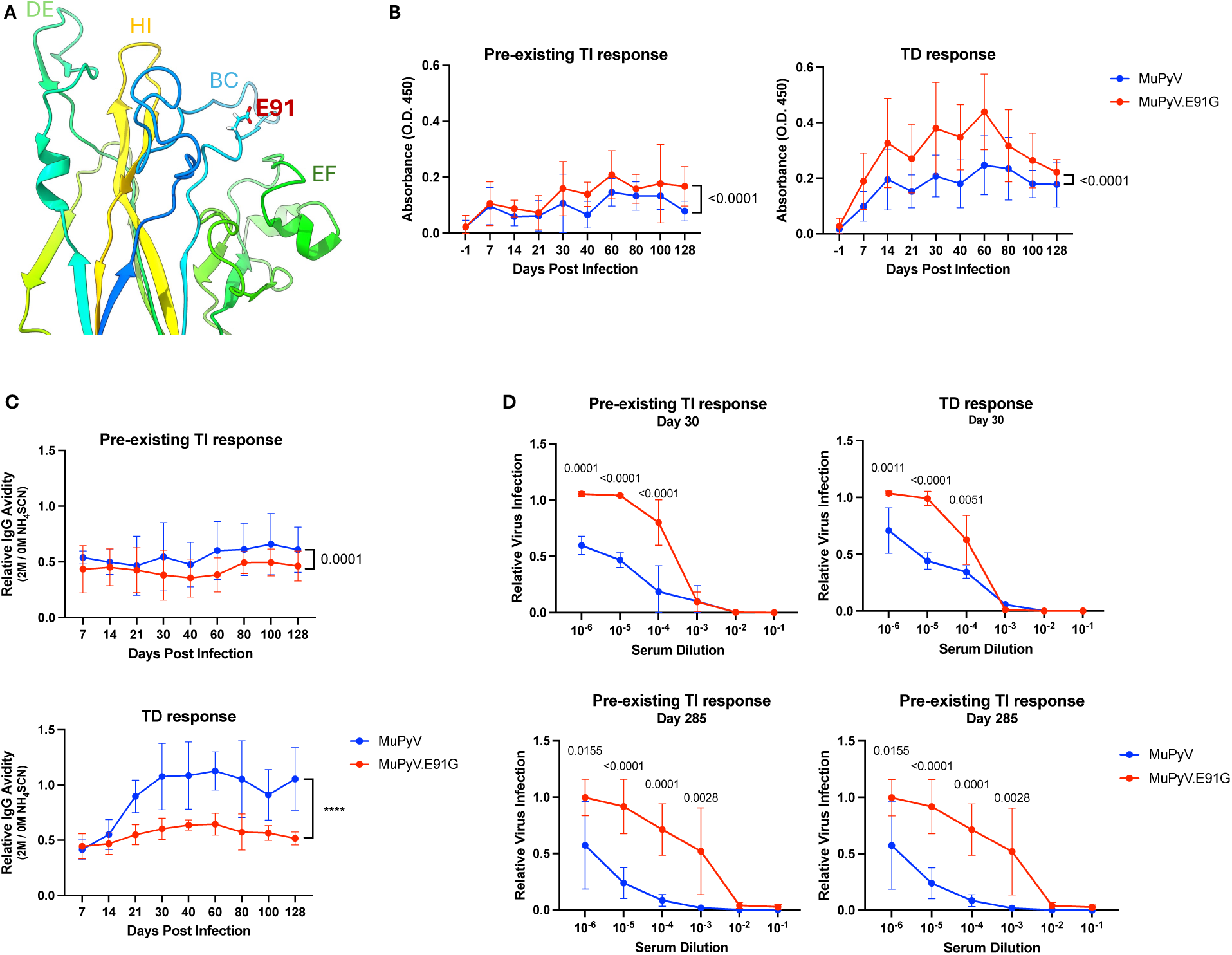
VP1 mutation E91G is an anti-viral IgG escape mutation. **(A)** Location of E91 residue in BC loop of VP1. (**B**) ELISA and (**C**) avidity using either MuPyV.E91G or MuPyV.WT as capture antigen. Sera from MuPyV.WT-infected mice (**left**) CD4 T cell depleted before infection or (**right**) given control rat IgG (n = 12). The absorbance at O.D. 450nm is graphed in (B). (**C)** Neutralization assays to MuPyV.WT or MuPyV.E91G. Sera from (**D, right)** pre-existing CD4 T cell-depleted mice or (**D, left**) control mice at (top) 30 or 285 dpi infected with MuPyV.WT (n = 8). Data are from 2 independent experiments. XY analysis non-linear regression fit with extra-sum-of-squares F test (**B-C**) and two-way ANOVA with Šídák’s multiple comparison test (**D**).

**Table 1:**
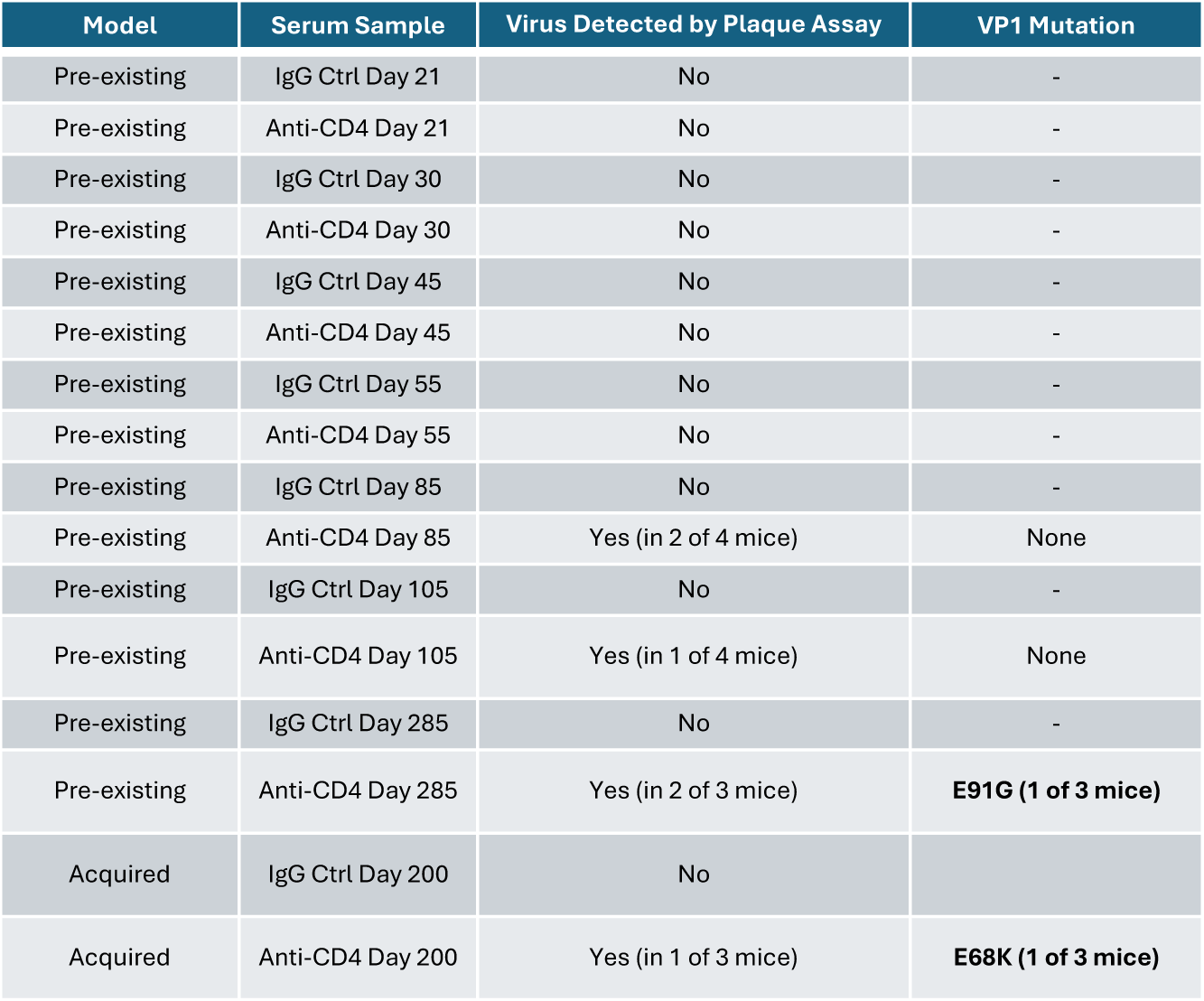
Outgrowth of VP1 mutant MuPyVs after serial passaging with sera from TI mice. Serum from CD4 T cell-depleted and rat IgG control-treated B6 mice was collected at the indicated timepoints after infection was serial passaged with MuPyV. After 4 serial passages in the presence of sera, infectious virus was detected by plaque assay, with plaques subjected to VP1 gene sequencing.

### Cryo-EM 2D image analysis of MuPyV-Fab complexes

Cryo-EM combined with single-particle analysis was used to visualize the MuPyV-Fab complex, where Fabs were from TI and TD mice sera. The 2D rotationally averaged images of control MuPyV particles from cryo-EM revealed circular profiles consistent with our previous study ^30^. Without Ab binding, the MuPyV particles displayed several concentric density layers within the capsid, corresponding to the encapsulated double-stranded viral DNA complexed with histone proteins (Fig. 5, orange region). The particle radius measured approximately 27 nm at the top surface of the capsomers (blue region, Fig. 5). 1D radial density profiles of control particle showed two prominent capsid peaks: one at 21 nm, corresponding to the capsid shell or floor, and another at 24 nm, representing the beta-jellyroll domain of the capsomer. In the Fab-bound particles, the capsid retained a similar radius of approximately 27 nm, excluding the additional densities that extended outward from the MuPyV surface (Fig. 5, blue region). The Fab density contributed two distinct peaks at radii of approximately 27 nm and 31 nm, representing the variable and constant regions of the Fab fragment, respectively.

**Figure 5.**
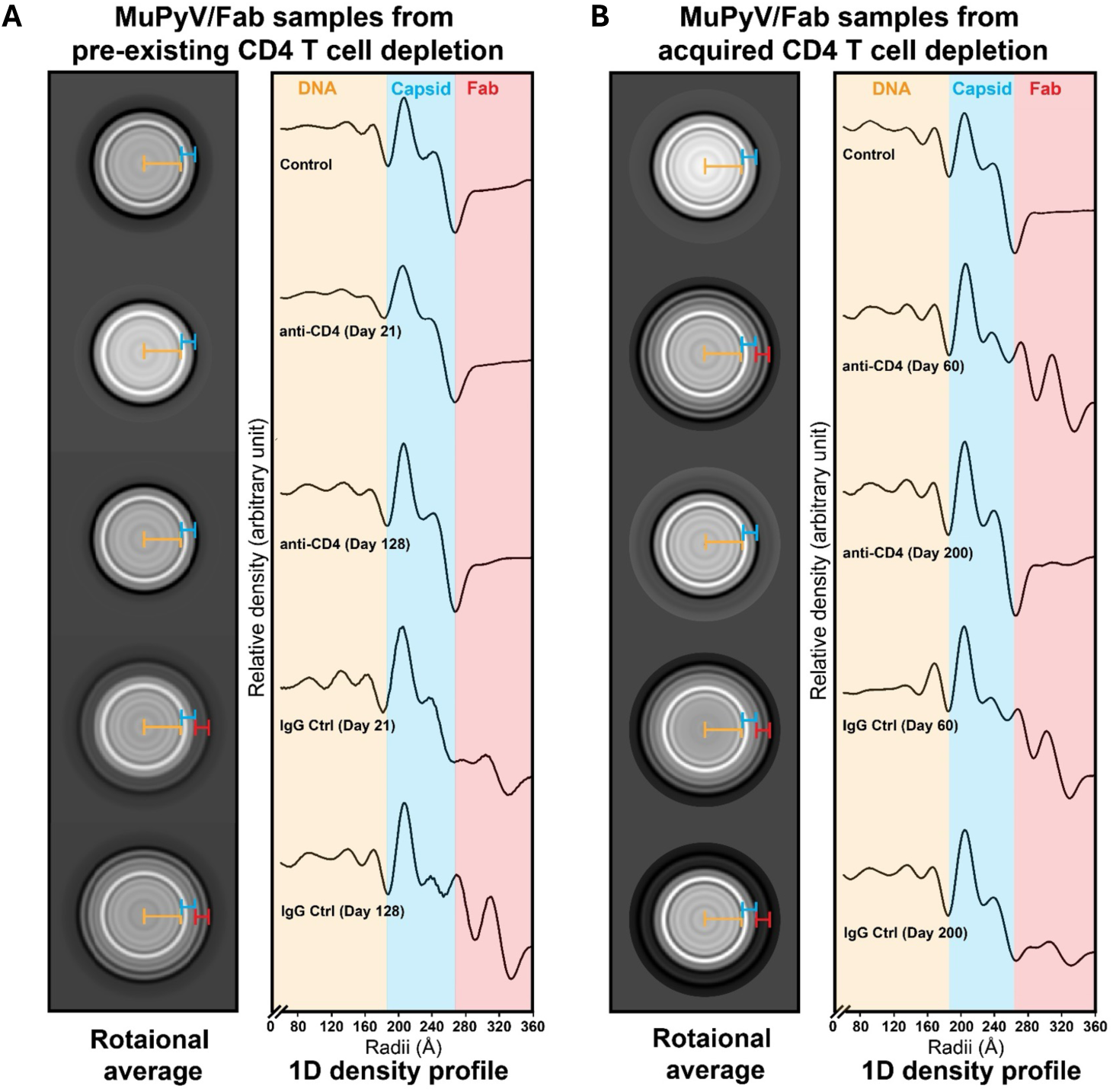
Radial density analysis of Fab binding to MuPyV capsid. Cryo-EM particles from each experimental condition were aligned, rotationally averaged, and analyzed for density distribution. (Left) Two-dimensional rotationally averaged images display concentric density rings, with protein density shown in white. The density gradient from high to low is indicated by grayscale shading. (Right) One-dimensional radial density profiles, derived from the 2D rotationally averaged images, illustrate Fab fragment binding on the MuPyV capsid surface. The X-axis represents the radius of the viral particle and the Y-axis shows the relative density distribution (unitless). Viral DNA, capsid, and Fab density regions are marked in orange, blue, and red, respectively. (A) Fab binding In mice with pre-existing CD4 T cell deficiency or control mice at 21- and 128-dpi. (B) Fab binding in mice with acquired CD4 T cell deficiency and control mice at 60- and 200-dpi. For cryo-EM experiments, only one mouse was selected from each experimental condition.

For pre-existing TI mice (Fig. 1A), polyclonal Abs (pAbs) showed no detectable Fab binding to the MuPyV capsid at either 21 dpi or 128 dpi (Fig. 5A). In contrast, Abs from TD mice demonstrated clear Fab binding to the capsid at both time points, with increased binding observed at 128 dpi, aligning with avidity results where TD Abs exhibited stronger binding at 128 dpi than at 21 dpi (**Fig. 1F**). In acquired TI mice (**Fig 3A**), pAbs displayed strong Fab binding to MuPyV capsid at 60 dpi, which diminished to undetectable levels by 200 dpi (**Fig. 5B**). In contrast, Abs from control mice exhibited consistent Fab binding at both 60 and 200 dpi. This data supports the idea of a transition from a TD to TI Ab response following acquired CD4 T cell loss (**Fig. 3E, 3H**). Overall, this cryo-EM data shows that TI Fabs bind poorly to MuPyV, consistent with the hypothesis that weak anti-VP1 IgG-virion capsid recognition is responsible for outgrowth of VP1 variants.

Despite effective virus neutralization by sera from pre-existing TI mice (**Fig. 1H**), no Fab density was observed in cryo-EM data. This discrepancy may stem from two factors: (1) purified Fab from these mice may bind at low occupancy per capsid particle, resulting in insufficient Fab density after averaging; or (2) effective neutralization may depend on Fc interactions, where even low concentrations of Ab could neutralize the virus through immune complex formation.

Individual mouse variability in antiviral Ab could also contribute to the observed differences in Fab density.

## 3D Reconstruction and Analysis of Ab Binding Sites on MuPyV Capsid

We generated 3D reconstructions using icosahedral averaging (**Fig. 6, Fig. S4, and Table S1**). In the Fab-free MuPyV capsid control, the structure displayed the expected T=7 icosahedral lattice, with 72 pentameric VP1 capsomers uniformly arranged across the viral surface. The innermost density layer corresponds to the histone-bound viral genome located beneath the capsid floor, while an additional density layer is linked to the extended C-terminal regions of VP1, which connect adjacent capsomers and reinforce capsid stability (**Fig. 6**).

**Figure 6.**
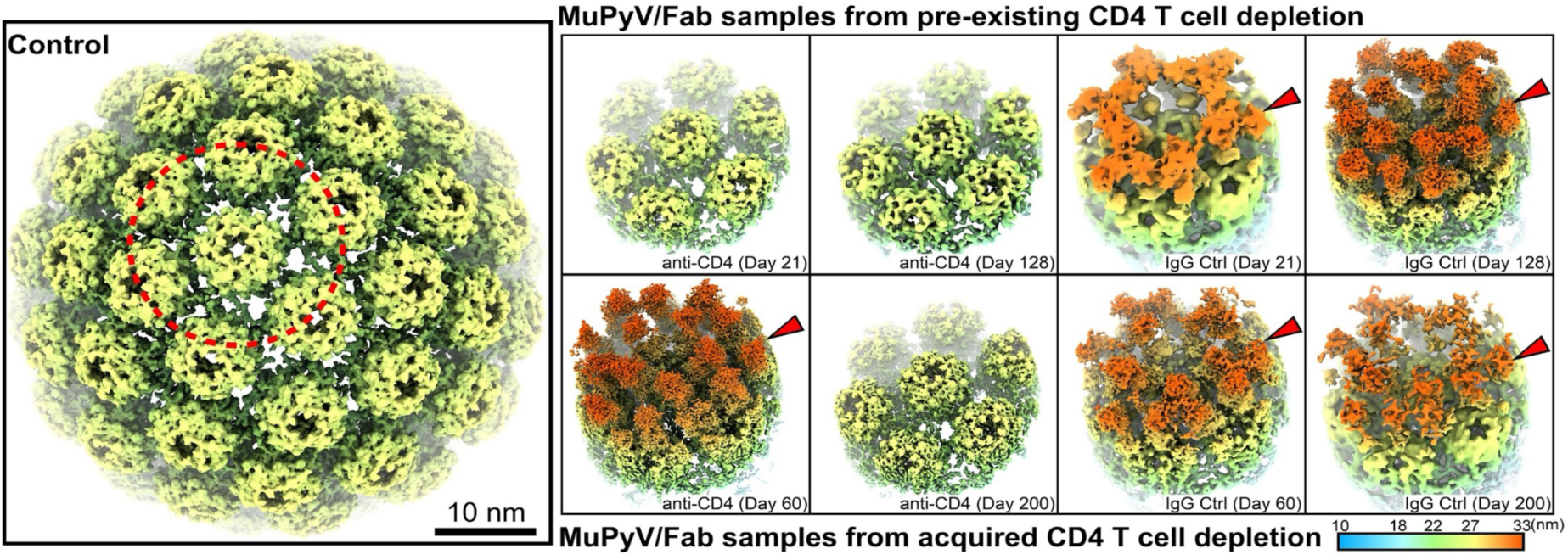
Cryo-EM 3D reconstruction of MuPyV in complex with anti-MuPyV Fabs under different experimental conditions. 3D reconstructions were generated with icosahedral symmetry imposed during data processing, and the resolution for each reconstruction is provided in Supplemental Table 1. Each MuPyV 3D reconstruction is displayed with a radial color scheme as indicated in the color legend, representing distances from the center in nanometers. (Left) A front half view of the control MuPyV capsid illustrates the capsomer organization, consistent with that of other polyomaviruses. (Right) Cut-away views of selected capsomers from hexameric pentamers highlight the density of bound Fab fragments on the MuPyV capsid surface. The upper panel represents Fab binding from pre-existing CD4 T cell deficiency conditions, while the lower panel shows results from the acquired CD4 T cell deficiency groups. Because the Fab fragments were derived from polyclonal Abs, the Fab density in each group reflects an average of different Ab types binding to the viral capsid.

In Fab-bound 3D reconstructions from pre-existing TI mice, no detectable Fab density was observed (**Fig. 6**). In contrast, TD Abs at 21 and 128 dpi displayed Fab densities clustered predominantly over the top surface of the VP1 capsomer (**Fig. 6**, red arrowheads). Each VP1 accommodated approximately one Fab density, concentrated around the BC- and HI-loops, with additional binding near the DE-loop on adjacent VP1 capsomers (**Fig. S5**), indicating that each Fab fragment may engage with antigenic loops spanning two neighboring VP1 capsomers. The specific residues involved in these interactions varied with Fab binding orientation. At 21 dpi, Fab density was tilted toward the quasi-threefold axis, producing a distinct smeared density pattern. In contrast, at 128 dpi Fab binding appeared more perpendicular to the VP1 top surface, an orientation difference highlighted in the radial color-coded density profiles (**Fig. 6**).

In reconstructions from acquired TI mice, Fab binding was evident at 60 dpi across multiple VP1 loops, including the DE-loop on adjacent VP1, consistent with 2D density observations (**Fig. 5**). At 200 dpi, Fab density diminished to undetectable levels, indicating a reduction in Fab affinity for VP1 binding over time as the Ab response transitions to TI IgG. In the TD mice, Fab densities were observed at both time points, although they appeared less densely packed at 200 dpi. Despite this reduced packing density, the Fab binding pattern remained relatively similar (**Fig. 6**).

In Fab-bound structures a thin density band extended along the capsomer surface, representing overlapping Fab regions within the icosahedral lattice of VP1. The orientations of bound Fabs varied, resulting in unique density patterns around capsid vertices. This flexibility in Fab binding angles and affinity could influence neutralizing efficacy and potentially lead to Fab occlusion effects during immune interactions. Despite differing binding patterns, VP1 residues involved in Ab interactions remained consistent across experimental conditions, suggesting that they are critical interaction sites in the anti-MuPyV humeral response (**Fig. S5**).

## DISCUSSION

In this study we explored how temporal differences in CD4 T cell deficiency during a persistent viral infection affected the nAb response and evolution of escape variants. We found that pre-existing and acquired loss of CD4 T cells in MuPyV-infected mice not only lowered virus-neutralizing IgG titers, but also reduced Ab avidity and clonal diversity, setting the stage for resurgent replication of WT virus and emergence of neutralization-resistant variants. By using intact MuPyV virions, we were able to directly extrapolate VP1 IgG functional analyses to cryo-EM reconstruction of VP1 Fab-virus binding. Unlike previous cryo-EM studies, including our own, which predominantly used mAbs for 3D reconstructions of Ab-virus complexes, this study employed Fabs from IgGs isolated from sera of infected mice ^30^. These findings support the use of cryo-EM to define virus-binding sites targeted by antiviral Abs in the sera of individuals with active infection or elicited by vaccination.

CSR and SMH can occur outside of the GC during TI Ab responses ^9,31,32^. We observed that MuPyV-specific IgM switched entirely to anti-viral IgG by 21dpi in TI mice, establishing that MuPyV infection induces CSR in the absence of GC reactions. TI IgGs, however, maintained poor avidity towards MuPyV, indicating a dissociation between CSR and SMH in the TI response to MuPyV. As detection of MuPyV-specific B cells is currently not technically possible, we compared BCR sequences of activated B cells (CD19^+^IgD^-^) from MuPyV-infected TI and TD mice. Although BCR diversity increased overtime in TD mice, there was no change in TI BCR diversity, confirming that CSR, but not SMH, occurs during TI Ab response during MuPyV persistent infection.

Despite GC reactions driving differentiation of B cells into LLPCs that reside in BM, LLPCs are also generated during GC-less TI Ab responses ^27,33,34^. We detected MuPyV-specific IgG-secreting ASCs in TI mice, even during late stages of persistent infection. This result led us to question if virus-specific ASCs generated during TI Ab response toward MuPyV were LLPCs. Administration of CD20 mAb has been shown to deplete GCs and memory B cells, but not affect established ASCs ^27^. We thus tested the longevity of the MuPyV TI ASCs with CD20 mAb-mediated B cell depletion given during persistent infection. At 70 dpi, both TI and TD mice given the anti-CD20 mAb had fewer MuPyV-specific IgG ASCs in the spleen than mice with their CD20 B cells intact (**Fig 2B**). This result indicates that depletion of CD20 B cells disrupted generation of new ASCs. In the BM, however, TD mice, but not TI mice, maintained MuPyV-specific ASCs after CD20 depletion, indicating that ASCs generated under TI conditions are not long-lived. Thus, maintenance of MuPyV-specific ASCs in TI mice depends on continuous generation of short-lived plasma cells. By extension, virus persistence may be required to re- supply short-lived TI ASCs to maintain TI anti-viral IgGs generated in the absence of GCs.

These results also raise a potential risk for loss of viral control for persistent infections in patients receiving CD20 mAb therapy for autoimmune diseases (e.g., multiple sclerosis) who have low or defective in CD4 T cells ^35^.

Unknowns in PML pathogenesis are: (1) the long latency between initiation of immune- modulating therapy and disease manifestations; and (2) the rarity of PML in at-risk populations given that most humans are JCPyV-seropositive ^36^. In the pre-antiretroviral therapy era, 5-8% of AIDS patients developed PML ^37^. Only 0.1% of MS patients receiving natalizumab infusions were diagnosed with PML, which was first detected after 24 months of receiving this a4 integrin blocking mAb ^38,39^. A growing number of chemotherapeutics and biologics garner FDA black box warnings of PML, including CD20 mAb therapies for MS and rheumatoid arthritis ^40,41^.

Retrospective analyses implicate absolute/relative CD4 T cell deficiency as the dominant immunological perturbation associated with PML. Giving a CD4 T cell depleting mAb to mice during the persistent stage of MuPyV infection models the “acquired” CD4 T cell deficiency antecedent for JCPyV-PML. With CD4 T cell depletion starting at 4 wk p.i., the anti-MuPyV IgG response mirrors that of immunocompetent infection-matched mice; approximately three months later, GCs are no longer detectable and anti-MuPyV IgG concentration and avidity progressively decline (**Fig 3**). Overall, these characteristics of the anti-MuPyV IgG response strikingly resemble the TI response in mice rendered CD4 T cell-deficient mice before infection. Together, these data support the conclusion that acquired CD4 T cell deficiency gradually shifts from a TD to a TI Ab response once the LLPCs fail to differentiate or survive during persistent polyomavirus infection. The extended timeframe for this TD-to-TI Ab shift following acquired CD4 T cell deficiency may be one factor contributing to the long latency for PML to develop in patients receiving natalizumab ^39^.

We recently demonstrated that passive immunization of B cell-deficient mice with a neutralizing rat VP1 mAb under TI conditions fostered outgrowth VP1 mutant escape variants in persistently infected mice ^1^. Serial passaging of MuPyV with TI immune sera showed a progressive loss in control of WT virus (3 mo p.i.) with a long latency to emergence of an E91G Ab-escape variant (10 mo p.i.) in only a fraction of mice (**Table 1**). Of note, E91 is a contact residue in the BC loop for our recent cryo-EM analysis of anti-VP1 rat mAb Fab-virus complexes [**Fig. S5D** and (*32*)]. The E91G VP1 mutation is an extensively characterized mutation that profoundly changes the profile of MuPyV-induced tumors, the magnitude of the host’s type I IFN response, and the efficiency of viral spread ^29,42^. Our data now show that this mutation also impairs neutralization by VP1-specific IgG. Using sera from acquired TI mice at 200 dpi for serially passaging MuPyV, we isolated additional viruses carrying the BC-loop mutation E68K (**Table 1**). Interestingly, the E68 is a contact residue conserved among VP1 Fabs isolated from several persistently infected TD mice as well as the rat VP1 mAb (**Fig. S5D**), suggesting that E91 and E68 are dominant amino acids in the BC loop for anti-VP1 IgGs.

Cryo-EM reconstruction of anti-VP1 Fab-virion complexes confirmed the functional loss VP1 epitope coverage. It is notable that most of the VP1 residues engaged by Fabs prepared from immune sera IgG overlapped with those we recently described using Fabs of a rat anti-VP1 mAb ^30^. Most of the common Fab contact residues were situated in the BC and HI loops, indicating surprisingly similar VP1 IgGs targets across two species. The ability to resolve individual VP1 contact points by immune sera IgG Fabs is possible only because of the highly restricted epitope range by the endogenous anti-VP1 IgG response. A further contraction of this already limited VP1 epitope repertoire and avidity due to CD4 T cell deficiency could open the door for outgrowth of VP1 nAb-escape viruses.

Our findings define the TI response in two clinically relevant models of PML: pre-existing and acquired CD4 T cell deficiency. In both models, TI conditions result in a quantitatively and qualitatively impaired VP1-specific IgG response that is conducive for outgrowth of nAb-escape viruses. Our findings raise four plausible insights into JCPyV-PML pathogenesis in CD4 T cell-deficient hosts: (1) the gradual shift from high- to low-avidity nAbs leading to breakthrough replication of WT virus in the kidney reservoir of persistent infection; (2) the stochastic nature of polyomavirus evolution such that Ab-escape variants emerge in only a few hosts; (3) the long timeframe for the TI antiviral IgG response to lose coverage of VP1 epitopes; and (4) the inter-host variation in the epitopes recognized by neutralizing IgGs. Together, these findings account for the years-long latency between start of immunomodulatory therapies and PML and the rarity of this devastating brain disease in at-risk individuals.

## MATERIALS AND METHODS

### Mice

C57BL/6 mice were purchased from the Jackson Laboratory (Bar Harbor, ME). B6.129-H2- *Ab1^tm1Gru^*N12 mice (MHCII KO) mice were purchased from Taconic Farms (Germantown, NY). B6.129-*IL21r^tm1wjl^*/Mmucd mice (IL21R KO) mice were purchased from the Mutant Mouse Resources and Research Centers at the University of California-Davis (Davis, CA). Male and female mice were 6-15 weeks of age. Same age mice were randomly assigned to experimental groups. Mice were housed and bred in accordance with the National Institutes of Health and AAALAC International Regulations. The Penn State College of Medicine Institutional Animal Care and Use Committee approved all experiments.

### Cell Lines and Primary Cells

NMuMG and BALB/3T3 clone A31 ‘A31’ were purchased from the ATCC. Cells were maintained in DMEM supplemented with 10% fetal bovine serum, 100 U/mL penicillin, and 100 U/mL streptomycin. Cell lines were authenticated by STR profiling (ATCC), confirmed to have the correct morphology, and were negative for mycoplasma.

### Virus Strains

All work was performed using the A2 strain of MuPyV. Viral stocks were generated by transfections of viral DNA into NMuMG cells with Lipofectamine 2000 Transfection Reagent (ThermoFisher). Viral amplification was done during a single passage of NMuMG cells. Virus stocks were titered on A31 fibroblasts by plaque assay ^43^.

### Generation of E91G MuPyV

The E91G mutation was introduced into the A2 MuPyV genome using the Quikchange II Site-directed mutagenesis kit (Agilent) as described ^29^.

### Virus Infections

Mice were infected with MuPyV s.c. in the hind footpad with 1 x 10^6^ PFU.

### *In vivo* Ab Administration and Flow Cytometry

CD4 T cells were depleted with rat anti-CD4 mAb (GK1.5). CD20 B cells were depleted with mouse anti-CD20 mAb (MB20-11, BioXCell). CD40L was blocked using Armenian hamster anti-CD40L mAb (MR1; BioXCell). Rat IgG, mouse IgG, and Armenian hamster IgG were used as controls, respectively. Cell depletion was confirmed in the peripheral blood every other week by collection from the superficial temporal vein and at euthanasia by staining with RM4-5 mAb (CD4) or RA3-6B2 mAb (B220) and 6D5 mAb (CD19) for CD4 T cell and B cell, respectively (ThermoFisher) (**Fig. S1A,C)**. GC B cells were defined as Live, CD19^+^, IgD^-^, GL7^+^, and CD95^+^ (**Fig. S1B**). Samples were run on a BD FACSSymphony17 flow cytometer (BD Biosciences) and analyzed using FlowJo software (Tree Star).

### Virus Purification

NMuMG cells were infected at a 0.1 multiplicity of infection (MOI) with MuPyV or MuPyV.E91G. Virus was purified from infected cell lysates and media as described ^30^.

### Viral Genome Quantification

Viral DNA was isolated from 50 μL of benzonase-treated purified virus and viral genomes quantified by TaqMan qPCR as described ^30^.

### ELISA and Avidity Assays

ELISAs were performed using 1×10^6^ genomes of purified MuPyV or MuPyV.E91G as capture antigen. Plates were treated with 1% BSA in 0.1% Tween PBS (blocking buffer). Mouse sera was diluted 1:150 in blocking buffer before being added to the plate. For avidity assays the virus:Ab complex was treated with 2M NH_4_SCN in 0.1 M phosphate for 15 minutes. Bound IgG in the ELISA and avidity assays was detected with an anti-mouse IgG specific secondary conjugated with HLP (Bethyl Laboratories), developed with 1-Step Ultra TMB-ELISA (ThermoFisher) and imaged using the Synergy HI plate reader with the absorption set at O.D. 450. IgG concentration was calculated using a standard curve of the VP1-specific rat mAb 8H7A5 (*32*) . IgG avidity at 2M NH_4_SCN was normalized to the absorption of samples without NH_4_SCN.

### ELISpot Assay

Plates were coated as described for the ELISA assays. Plates were blocked with 5% BSA in 0.1% Tween PBS (blocking buffer). Single cell suspensions of spleen and BM were treated with ACK. Lymphocytes were isolated from the kidney by digestion with collagenase followed by centrifugation on a 44%/66% Percoll gradient (ref). Cells were added to the ELISpot plates at a serial dilution of 1:5 in DMEM with 8% FBS and 2% EDTA starting with 1×10^5^ cells/well.

Anti-mouse IgG conjugated with biotin (Mabtech) was added at a 1:2000 dilution. Streptavidin – ALP (Mabtech) was added at 1:2000 dilution. Plates were developed using BCIP/NBT-plus for ALP reagents (Mabtech) and read on ImmunoSpot Analyzer (Cellular Technology Limited). Data shown is the total number of ASCs per organ.

### Immunofluorescence Microscopy

Spleens were fresh-frozen in Tissue-Tek O.C.T. Compound (Sakura) on dry ice prior to cryosectioning. Sections were fixed to the slide with 4% PFA and stained with anti-GL7, anti- IgD, and anti-CD4 antibodies. Samples were mounted with Prolong Gold Antifade Mountant with DAPI (ThermoFisher). Slides were imaged on a Leica DM4000 fluorescence microscope in blinded fashion. GCs was counted for the entire spleen section. Adjustments for brightness/contrast were done uniformly to all images in the group using LAS X (Leica). GC area was calculated by Image J.

### BCR Sequencing

TI and TD mice spleens were harvest at 21 and 128 dpi and put into single cell suspension. Cells were stained and sorted on the BD FACSMelody cell sorter for live, single-cell, IgD^-^CD19^+^ B cells. RNA was isolated from sorted B cells using Invitrogen Purelink Viral RNA/DNA Mini Kit (ThermoFisher Scientific) and sent to Cellecta for BCR sequencing using their DriverMap Adaptive Immune Receptor (AIR) profiling assay. Samples were amplified, checked for QC quality, and incubated with a mix of reverse mouse AIR mouse BCR Gene-specific (GS) primers. The resulting RNA-RevGSP product was purified, and cDNA was made. cDNA was extended by incubation with master mix containing Forward FR3 AIR mouse GSPs. Purified cDNA product was quantified by Qubit fluorescence measurement and underwent next-gen sequencing using NextSeq500. Bioinformatics was also performed by Cellecta. MiXCR was used to align the sequencing reads and identify clonotypes and their abundances ^44^.

### Infection Neutralization Assay

Sera from CD4 T cell-depleted and IgG control mice was diluted 10-fold from 1:10 to 1:1,000,000 and incubated with MuPyV of MuPyV.E91G at 4°C. The virus:sera mixture was added to NMuMG cells and incubated on ice for 1.5 h then for 24 h at 37°C. mRNA was harvested with TRIzol Reagent (ThermoFisher) and isolated by phenol: chloroform extraction by isopropanol precipitation. cDNA was prepared with random hexamer and Revertaid RT (ThermoFisher). LT mRNA levels were quantified by Taqman qPCR and normalized to TATA-Box Binding Protein (IDT) ^45^. Fold expression (2^-(ΛΛCt)^) was calculated against virus not incubated with sera.

### Viral DNA Isolation and Quantification

Tissue of interest was homogenized using a TissueLyser II (Qiagen). The Wizard Genomic DNA Purification Kit (Promega) was used to isolated DNA that was then quantified by qPCR. Quantification of viral DNA was calculated based on a standard curve.

### Sera-mediated Selection of VP1 Mutant Viruses

NMuMG cells were infected with MuPyV at a MOI or 0.1. Sera from TI and TD mice were normalized based on ELISA to contain the same concentration of anti-MuPyV IgG. 100 ug MuPyV-specific IgG was added 24 h p.i. New sera was added when media was changed every 5 d. Lysates were collected every 1.5 wk when cell death was observed. Virus was isolated from the lysate, diluted 1:100, and added with sera to NMuMG cells. After 4 passages, the final lysate was collected and subjected to plaque assay to determine the presence of virus. Viral DNA in the lysate was isolated using the Pure-Link Viral RNA/DNA mini Kit (ThermoFisher) if plaques were observed. The VP1 region of the genome was amplified by PCR and sequenced. Identified mutations were cloned and generated by site-directed mutagenesis.

### Fab Isolation for Cryo-EM

Fabs were isolated from sera of CD4 T cell depleted and IgG control mice using the Pierce Fab Micro Preparation Kit (Thermo Fisher) ^30^.

### Sample Vitrification, Cryo-EM Data Collection, and Image Processing

Purified Fab samples were mixed with MuPyV at molar ratios of Fab to VP1 subunits exceeding 1:1. The mixtures were incubated on ice for 1h and then concentrated using 100K Omega Nanosep filters in a HERMLE Benchmark Z216-MK refrigerated microcentrifuge. Samples were centrifuged at 1,500 g at 4°C until 20-40 μL of concentrated sample was obtained for vitrification. For each grid preparation, 4 μL of the sample mixture was applied to glow-discharged grids, either ultrathin continuous carbon film-coated (CF300-Cu-UL, EMS) or Quantifoil R2/2 holey carbon-film coated copper grids. The grids were vitrified using a Thermo Fisher Scientific Vitrobot Mark IV with blotting parameters set to 4°C, 95–100% humidity, a blotting force of 0, a blotting time of 4 seconds, and a wait time of 10 s before plunge-freezing into liquid ethane cooled to -180°C by liquid nitrogen.

The frozen-hydrated grids were clipped into autogrids and loaded into a Thermo Fisher Scientific Krios G3i electron microscope, operated at 300 kV under liquid nitrogen temperature.

Data were collected at varying magnifications depending on the observed sample concentration (**Table S1**). Data acquisition was performed using EPU automation software, with each image recorded as a series of movie frames at a total dose of 30 electrons/Å² and a frame dose rate of approximately 1 electron/Å². A Gatan Bioquantum energy filter with a 30 eV slit was used, and all images were collected at defocus values between -1 and -3 μm on a K3 camera using super-resolution counting mode. Image frames were transferred to a high-performance computing cluster for further processing.

Cryo-EM data processing was conducted using Relion (v3.1) and EMAN2 (v2.91) ^46,47^.

Movie frames were aligned and corrected for beam-induced motion using MotionCor2 in Relion, after which the images were binned by 2 to enhance the signal-to-noise ratio. Particle picking was initially performed on a subset of data, either manually or semi-automatically using the Laplacian-of-Gaussian filter. After generating 2D class averages, these averages were used as templates for further automated particle picking across all micrographs. Extracted particles underwent reference-free 2D classification in Relion, and particles with contaminated ice or poor quality were discarded. For 2D image analysis, rotational averages and 1D density profiles were generated in EMAN2. For 3D reconstruction, initial models were created de novo in Relion, and all datasets displayed the characteristic T=7 capsomer arrangement of polyomaviruses. These models were further refined in Relion, with final resolution estimated by gold-standard Fourier shell correlation (Table S1).

For visualization, surface-shaded 3D models were rendered using UCSF ChimeraX ^48,49^.

Atomic model building was based on the highest-resolution cryo-EM map with PDB: 7K22 serving as the starting structure. The model was iteratively refined in ISOLDE, real-space refined in PHENIX, and final adjustments were made in Coot following previously established protocols^50–53^. Model quality was assessed using MolProbity ^54^. This final model was used as a reference for analyzing Ab-interacting residues in UCSF ChimeraX.

### Statistical Analysis

Prism 8 software (GraphPad) was used for statistical analysis. Tests included Two-way ANOVA with Tukey’s or Šídák’s multiple comparisons test, XY analysis non-linear regression fit with extra-sum-of-squares F test, and student T test. P values of <0.05 were considered significant and significant differences were labeled. All data are shown as mean with error bars representing +/- standard deviation (SD). Statistical methods were not used to pre-determine sample size. Figures contain the data from all repeats and no data points were excluded. All sample sizes, number of repeats, and statistical tests are included in the Figure Legends.

## Data Availability

All maps and models generated by Cryo-EM will be deposited to wwPDB.

## Supplemental Materials

Figure S1: Flow cytometry gating for confirming cell depletions and GC B cells

Figure S2: Mice with an impaired CD4 T cell compartment recapitulates the CD4-depleted MuPyV-specific TI IgG response.

Figure S3: CD40L blockade resembles the impaired anti-viral IgG response in acquired CD4 T cell deficiency.

Figure S4: Radially color-coded cryo-EM reconstructions of MuPyV capsids with anti-MuPyV Fab binding under various conditions.

Figure S5. Image analysis of MuPyV VP1 residues interacting with various anti-MuPyV Fabs. Table S1. Cryo-EM data collection and image process statistics.

## ACKNOWLEDGEMENTS

We thank the staff of the Penn State College of Medicine Flow Cytometry Core Facility (RRID: SCR_021134) and the Comparative Medicine Histology Core. We thank the Penn State College of Medicine for access to TEM (RRID:SCR_021200), cryo-EM (RRID: SCR_021200) and the HPC (RRID:SCR_022953) core facilities. We thank the members of the Lukacher and Wang laboratories for their valuable discussion and feedback of this study.

## AUTHOR CONTRIBUTIONS

Conceptualization: DJG, AEL, SLH, JCYW, KNA, MDL Methodology: KNA, MDL, KMA, GJ, KP, AO, SG

Investigation: KNA, MDL, KMA, GJ, AO, SG Visualization: KNA, JCYW

Funding acquisition: AEL, JCYW, SLH, KNA Project administration: KNA, AEL, JCYW Supervision: SJE, MJM, JLS, EH

Writing – original draft: KNA, JCYW Writing – review & editing: AEL

## DATA AND MATERIALS AVAILABILITY

All data are available in the main text or the supplementary materials.

