## Supplemental Figs 1-5 & Supplemental Table 1 for "The CD4 T cell-independent IgG response during persistent virus infection favors emergence of neutralization-escape variants"

### Supplemental Materials

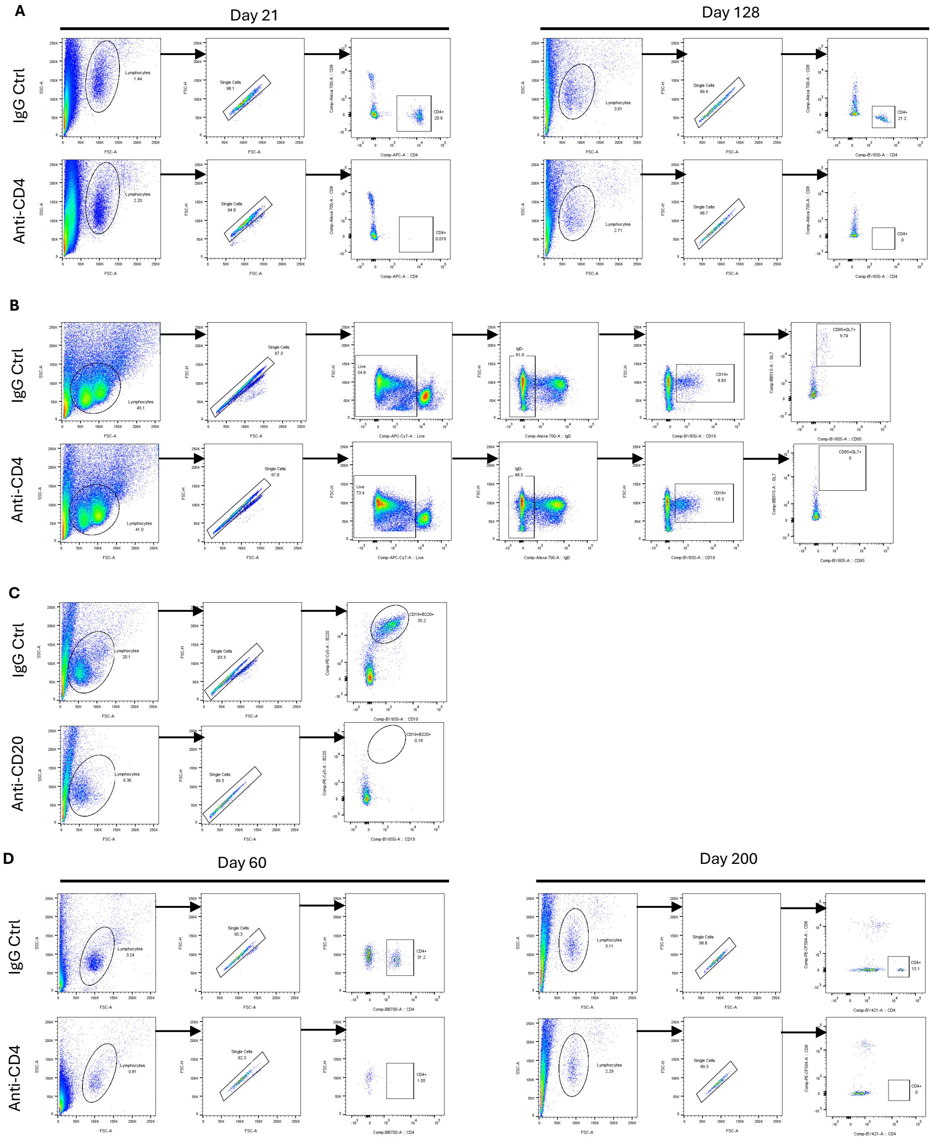


**Supplemental Figure 1: Flow cytometry gating for confirming cell depletions and GC B cells.** Blood was collected via submandibular vein puncture every two weeks and test via flow cytometry for A, C, and D. All shown data is representative. **(A)** CD4 T cell depletion in mice given GK1.5 mAb prior to infection at 21 and 128 dpi. Samples were gated for lymphocytes, single cells, and CD4 T cells. **(B)** GC B cells in the spleen as gated by lymphocytes, single cells, live cells then IgD$^-$ CD19$^+$ CD95$^+$ GL7$^+$. **(C)** CD20 B cell depletion as determined by gating for lymphocytes, single cell, and CD19$^+$ B220$^+$ B cells. **(D)** CD4 T cell depletion starting during persistent infection (at 28 dpi). Gating strategy was the same as in **(A)**.

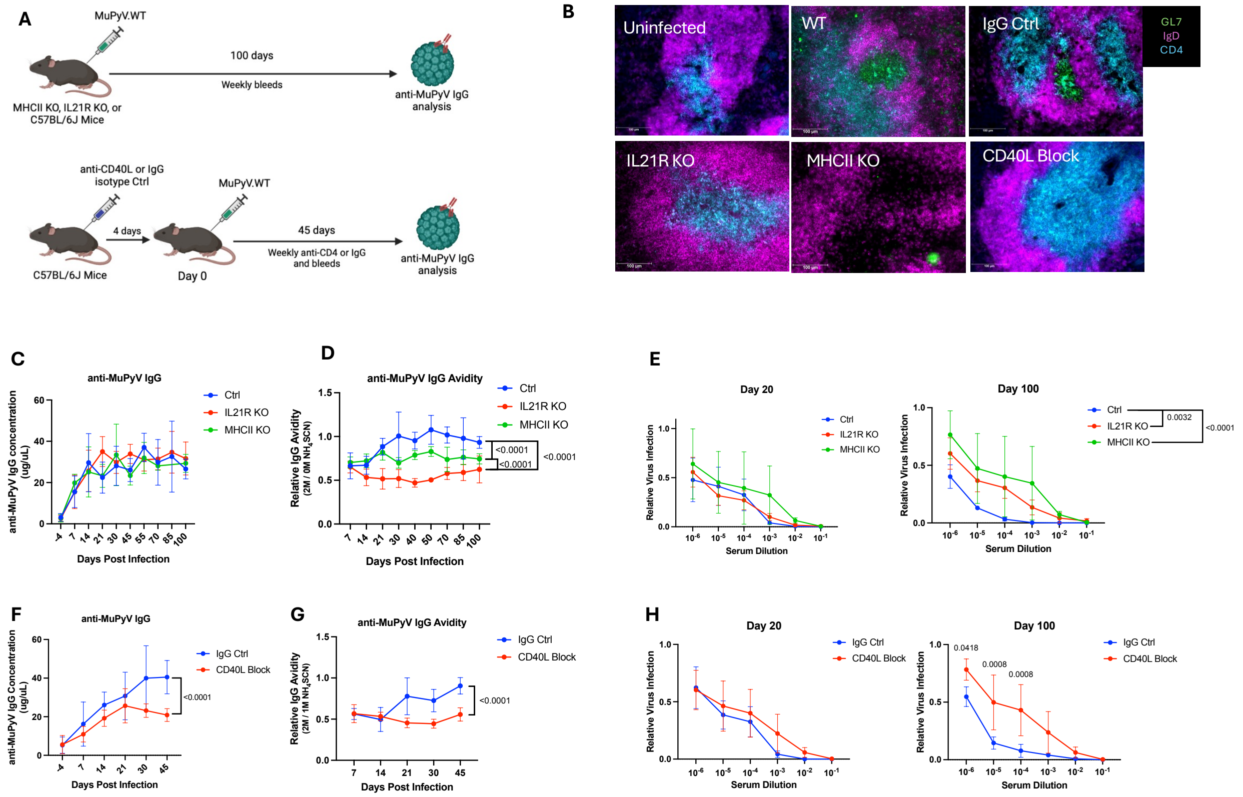


**Supplemental Figure 2: Mice with an impaired CD4 T cell compartment recapitulates the CD4-depleted MuPyV-specific TI IgG response. (A)** Experimental setups **(Top)** MHCII KO, IL21R KO, and B6 mice were infected S.C. with MuPyV. Image generated with Biorender. **(Bottom)** B6 mice were given CD40L blocking mAb or Armenian hamster IgG control at -4 and -1 dpi, then weekly after virus inoculation. **(B)** Representative IF images of GCs stained for GL7 (green), IgD (magenta), and CD4 (cyan) from spleens at 20 dpi (n = 4). MuPyV-specific IgG concentration by ELISA of **(C)** MHCII KO and IL21R KO sera and **(F)** CD40L (n = 8-12). Avidity towards MuPyV from **(D)** IL21R KO and MHCII KO and **(G)** CD40L blockade sera (n = 8-12). Neutralization of MuPyV by serum from **(E)** IL21R KO and MHCII KO or **(H)** CD40L blockade sera from **(left)** 20 dpi or **(right)** 100 dpi. Error bars are mean ± SD. Data are from 2-3 independent experiments. Data was analyzed by XY analysis non-linear regression fit with extra-sum-of-squares F test (**C-D, F-G**); and two-way ANOVA with Šídák's multiple comparison test (**E, H**).

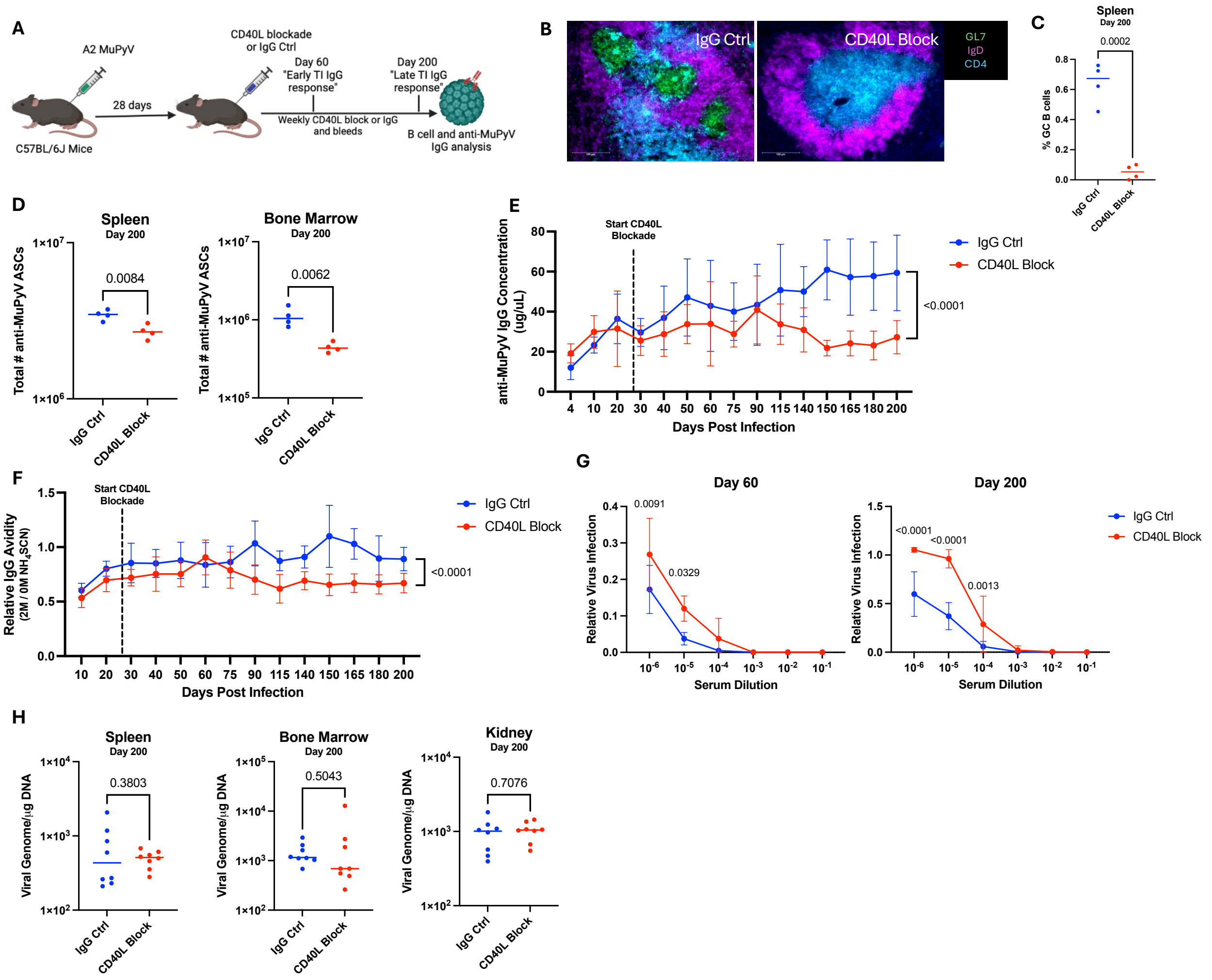


**Supplemental Figure 3: CD40L blockade resembles the impaired anti-viral IgG response in acquired CD4 T cell deficiency. (A)** Experimental setup. B6 mice infected with MuPyV received a CD40L blocking mAb or Armenian hamster IgG control weekly starting at 28 and 30 dpi. Image generated with Biorender. **(B)** Representative IF images of GCs [GL7 (green), IgD (magenta), and CD4 (cyan)] CD40L blockade or control IgG mouse spleens at 200 dpi (n = 4). **(C)** Frequency of B220$^{+}$ CD19$^{+}$ IgD$^{-}$ GL7$^{+}$ CD95$^{+}$ GC B cells in the spleen of CD40L blockade or IgG control mice at 200 dpi (n = 4). **(D)** ELISpot assays of MuPyV-specific ASCs in the **(left)** BM or **(right)** spleen (n = 4). **(E)** ELISA and **(F)** avidity assays using purified MuPyV as capture antigen (n = 8). **(D)**. Avidity in presence of 2M $NH_4SCN$ was normalized to samples treated with 0 M $NH_4SCN$ **(F)**. (**G**) Neutralization of MuPyV by serum taken at 60 dpi **(left)** or 200 dpi **(right)** (n = 4). (**H**) Viral DNA levels in the BM, spleen, and kidney at 200 dpi (n = 7-9). Error bars are mean ± SD from 1-2 independent experiments. Data was analyzed by Student's *t*-test (**C, D, H**); XY analysis non-linear regression fit with extra-sum-of-squares F test (**E, F**); and two-way ANOVA with Šídák's multiple comparison test (**G**).

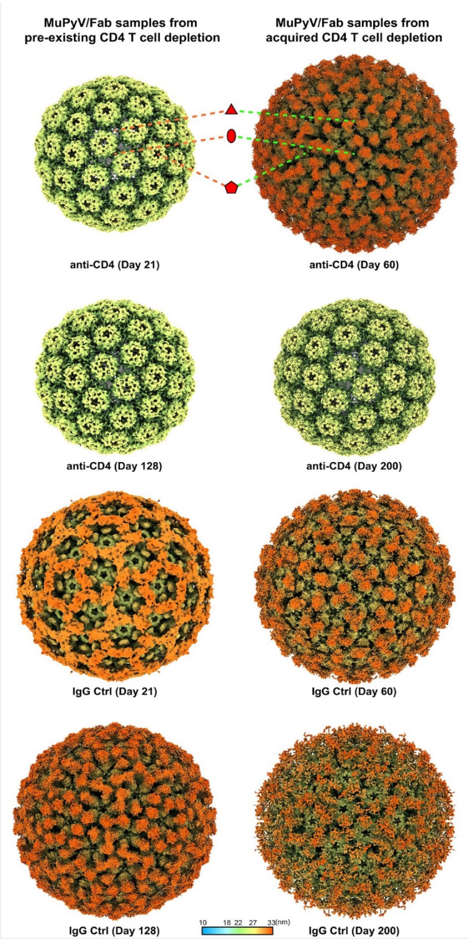


**Supplemental Figure 4: Radially color-coded cryo-EM reconstructions of MuPyV capsids with anti-MuPyV Fab binding under various conditions.** Surface-shaded representations of MuPyV capsids bound to anti-MuPyV Abs are shown along the icosahedral twofold axis. The 3D reconstructions are radially color-coded according to the scale bar at the bottom. All maps without Fab densities display a consistent external appearance, characterized by 72 pentameric capsomers arranged in a T=7 icosahedral lattice, with radii ranging from approximately 19 to 27 nm, colored from green to yellow. For maps exhibiting Fab densities, the Fab fragments bind to the top surface of VP1 capsomers, extending up to 34 nm in radius. Due to the polyclonal nature of the Abs, multiple Fab types are present, and the resulting structures are products of icosahedral averaging. All Fabs consistently bind at the top surface of the VP1 capsomers. The Fab fragments orientation varies slightly across different binding angles, as shown in the 3D reconstructions, with some regions exhibiting subtle differences in Fab density. Oval, triangle, and pentagon symbols indicate the locations of icosahedral twofold, threefold, and fivefold symmetry axes, respectively. Scale bar: 23 nm.

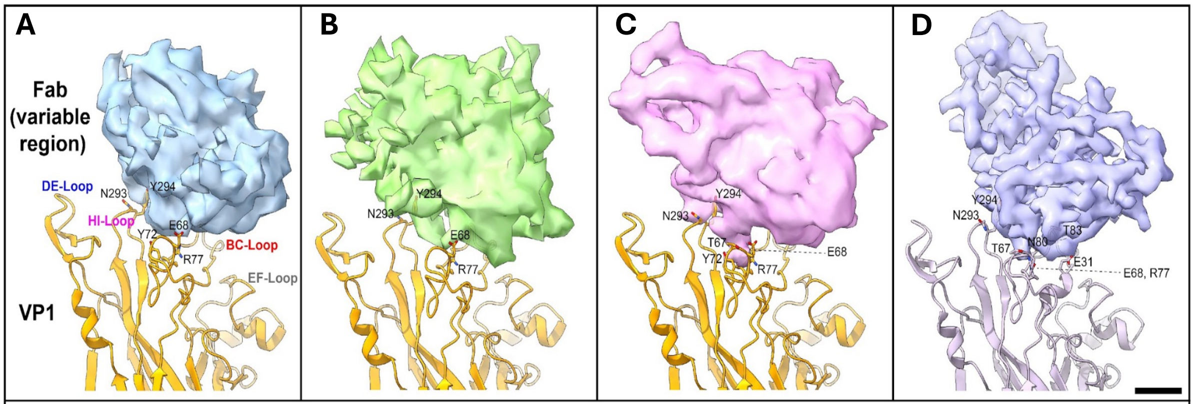


| | | | |
|---|---|---|---|
| **Residues of VP1 in close proximity to current Fab binding** | | | |
| Glu68, Gly69, Tyr72, Arg77, Gly78, Asn293, Tyr294 | Glu68, Gly69, Arg77, Gly78 Asn293, Tyr294 | Thr67, Glu68, Tyr72, Arg77, Gly78, Asn293, Tyr294 | Glu68, Gly69, Arg77, Asn80, Thr83, Glu91, Asn293, Tyr294 |
| **Residues of VP1 in close proximity to neighboring Fab binding** | | | |
| Asn149 | Asp85 | Asp85, Asn149 | Phe141, Lys151 |

**Supplemental Figure 5. Image analysis of MuPyV VP1 residues interacting with various anti-MuPyV Fabs.** Schematic representation of Fab fragment binding to VP1, highlighting the residues involved in Ab interactions. Each panel shows a cryo-EM density map of the Fab bound to a VP1 subunit from a pentamer, with modeled atomic structures illustrating key interacting residues. The variable region of the Fab density was segmented to visualize the specific loops involved in binding. Residues were considered interacting if their side-chain atoms were within a 4 Å radius, as determined by van der Waals contact criteria. We focused our analysis on 3 structures **(A, B**, and **C)** with the most distinct Fab density. **(A)** Polyclonal Ab taken at 128 dpi from control mice given rat IgG starting prior to infection. **(B)** Polyclonal Ab collected at 60 dpi from a mouse given IgG control starting at 28 dpi. **(C)** Polyclonal Ab from an animal with acquired CD4 T cell deficiency at 60 dpi. **(D)** The neutralizing rat anti-MuPyV VP1 mAb 8A7H5 (PDB: 7K22). The table below each panel lists specific VP1 residues in close proximity to the current Fab binding (highlighted in red, magenta, and blue for each loop) and residues accessible for neighboring Fab interactions (including key residues such as Asp85, Asn149, Phe141, and Lys151). Scale bar: 1 nm.

| 3D reconstructions from pre-existing CD4 T cell depletion | Cont rol | anti-CD4 (Day 21) | anti-CD4 (Day 128) | IgG Ctrl (Day 21) | IgG Ctrl (Day 128) |
|---|---|---|---|---|---|
| **Resolution (Å)** | 5 | 6.7 | 4.6 | 7.3 | 5.6 |
| **Pixel size (Å)** | 2.16 | 2.16 | 1.4 | 2.16 | 2.16 |
| **3D: particle #** | 8701 | 3050 | 18863 | 3132 | 6865 |
| **3D reconstructions from acquired CD4 T cell depletion** | **-** | **anti-CD4 (Day 60)** | **anti-CD4 (Day 200)** | **IgG Ctrl (Day 60)** | **IgG Ctrl (Day 200)** |
| **Resolution (Å)** | - | 4.3 | 3.7 | 6.6 | 9.3 |
| **Pixel size (Å)** | - | 1.4 | 1.4 | 2.68 | 2.8 |
| **3D: particle #** | - | 18802 | 86178 | 6633 | 2970 |

**Supplemental Table 1. Cryo-EM data collection and image process statistics.**
